# Phenolic profiles and antioxidant activities of Saskatchewan (Canada) bred haskap (*Lonicera caerulea)* berries

**DOI:** 10.1101/2021.08.04.455127

**Authors:** Lily R. Zehfus, Christopher Eskiw, Nicholas H. Low

## Abstract

Phenolic extracts from five Saskatoon, Saskatchewan, bred and grown haskap berry varieties (Aurora, Blizzard, Honey Bee, Indigo Gem, and Tundra) were characterized via liquid chromatography with photodiode array detection (HPLC-PDA) and mass spectrometry (HPLC-MS/MS). Tundra had the highest phenolic content (727.0 mg/100 g FW) while Indigo Gem had the highest anthocyanin content (447.8 mg/100 g FW). HPLC-MS/MS identified two previously unreported anthocyanins (Tundra variety): delphinidin-sambubioside and a peonidin-pentoside. Fruit extracts were fractionated to produce an anthocyanin rich (40% ethanol) and flavanol/flavonol rich (100% ethanol) fraction. This process affords the ability to isolate/concentrate specific subclasses for nutraceutical applications. High *in vitro* radical scavenging was observed for all haskap phenolic extracts. An extract from the Tundra variety delayed borage oil oxidation more effectively than commercial antioxidants (BHT and Rosamox). These results show the high phenolic content of these haskaps along with their capacity for radical scavenging/delaying lipid oxidation, indicating potential commercial value.

## 1. Introduction

Haskap berries (*Lonicera caerulea*) are an edible blue honeysuckle species that grow wild in northern boreal climates of the Americas and Eurasia. This species has been an integral part of domesticated horticulture in parts of Japan (e.g. the Kurile Islands) and Russia for over a century. Haskaps began to gain interest in North America in the 1990s as data regarding their phenolic content became more widely available (Celli, Ghanem, & Brooks, 2014). Like many other nontraditional berry species (e.g. lingonberries), haskaps have been found to have much higher phenolic content than some more common berries (e.g. blueberries). Phenolics are secondary metabolites in plants with a variety of functions including antimicrobial activity and UV protection (Pandey & Rizvi, 2009; Bhattacharya, Sood, & Citovsky, 2010; Cheynier, 2012; Giada, 2013); however, the primary interest in phenolics arises from their free radical scavenging (RS) abilities which can reduce oxidative stress in living organisms.

Haskap berry phenolic research has mainly focused on Russian, Polish, and Japanese varieties with the most abundant phenolic subclass as anthocyanins which account for the berries’ intense purple pigmentation (Chaovanalikit, Thompson, & Wrolstad, 2004; Kusznierewicz, Piekarska, Mrugalska, Konieczka, Namiesnik, & Bartoszek, 2012; Caprioli et al., 2016). This subclass, along with other members of the flavonoid class (e.g. flavanols and flavonols), are often reported to be very effective free radical scavengers in comparison to other phenolics (Rice-Evans, Miller, & Paganga, 1996; Kähkönen & Heinonen, 2003; Wang, Chen, Sciarappa, Wang, & Camp, 2008). As such, haskap berries have potential as a dietary source of nutraceutical compounds to improve health. Select haskap phenolic extracts have demonstrated possible health benefits including: (a) the inhibition of pro-inflammatory cytokines (e.g. interleukin-6 and tumor necrosis factor-α) in human monocytes (Rupasinghe, Boehm, Sekhon-Loodu, Parmar, Bors, & Jamieson, 2015); (b) decreased expression of adipogenic gene transcripts in hepatic carcinoma cancer (HepG2) cells (Park, Yoo, Lee, & Lee, 2019); and (c) improved measurements of working memory and lower diastolic blood pressure and heart rate in adult males (Bell & Williams, 2018). In addition to health benefits, the food industry could utilize haskap berry phenolic extracts as they have the potential to extend the shelf-life of food products by reducing the negative impacts of lipid oxidation on the colour, flavour, odour, nutritional quality, texture, and appearance of foods.

Interest in haskap berries in North America led to the establishment of a Canadian breeding program at the University of Saskatchewan’s Horticulture Field Lab, which resulted in the introduction of a selection of recently developed varieties available to the public. Much less is known about the phenolic composition and antioxidant potential of these varieties. It was hypothesized that Saskatchewan grown and bred haskaps would have high anthocyanin and overall phenolic content with significant *in vitro* radical scavenging capabilities. Also, the application of solid phase extraction (SPE) could be used to produce fractions with different antioxidant potentials based on their phenolic composition/structures, which could find use in commercial applications. To address these hypotheses, the following research goals were investigated: (a) determine the phenolic composition of five Saskatchewan-grown and bred haskap berry varieties (Aurora, Blizzard, Honey Bee, Indigo Gem, and Tundra) using high performance liquid chromatography (HPLC) and HPLC tandem mass spectrometry (HPLC-MS/MS) and assess their *in vitro* free radical scavenging activities; (b) SPE fractionation of haskap berry extracts coupled with phenolic structural analysis and *in vitro* free radical scavenging activities of the produced fractions; and (c) assess the antioxidant potential of the Tundra haskap phenolic rich extract when added to borage oil via rancimat analysis.

## 2. Materials & methods

### 2.1 Samples

Aurora, Blizzard, Honey Bee, Indigo Gem, and Tundra haskap berry varieties (2017 crop) were obtained from the University of Saskatchewan Horticulture Field Laboratory (Saskatoon, SK, Canada). Berries were stored at −30 ± 2°C until analysis. Borage oil was obtained from Bioriginal Food & Science Corporation (Saskatoon, SK, Canada) and commercial rosemary extract (Rosamox) was obtained from Kemin Industries Inc. (Des Moines, IA, USA).

### 2.2 Chemicals

The following chemicals were obtained from Sigma-Aldrich Canada Ltd. (Oakville, ON): Amberlite^®^ XAD16N resin; arbutin; 2,2’-azinobis(3-ethylbenzothiazoline-6-sulfonic acid) (ABTS); caffeic acid; catechin; chlorogenic acid; *p-*coumaric acid; cyanidin chloride, cyanidin-3-*O*-rutinoside; cyanidin-3,5-*O*-diglucoside; 2,2’-diphenyl-1-picrylhydrazyl (DPPH); epicatechin; ferulic acid; gallic acid; 4-hydroxybenzoic acid; naringenin; phloridzin; potassium persulfate; quercetin; and rutin (quercetin-3-*O*-rutinoside).

The following chemicals were purchased from Santa Cruz Biotechnology (Dallas, TX, USA): 6-hydroxy-2,5,7,8-tetramethylchroman-2-carboxylic acid (Trolox) and vanillic acid. Cyanidin-3-*O*-xyloside was obtained from Toronto Research Chemicals Inc. (Toronto, ON). Apigenin, cyanidin-3-*O*-arabinoside, cyanidin-3-*O*-glucoside, isorhamnetin-3-*O*-rutinoside, pelargonidin-3-*O*-glucoside, peonidin-3-*O*-glucoside, and quercetin-3-*O*-galactoside were purchased from Extrasynthese S. A. (Genay, France). Quercetin-3-*O*-glucoside and isorhamnetin-3-*O*-glucoside were obtained from Indofine Chemical Company, Inc. (Hillsborough, NJ, USA).

Chemicals obtained from Fisher Scientific (Edmonton, AB) were: acetonitrile; butylated hydroxytoluene (BHT); formic acid; methanol; phosphoric acid; sodium carbonate; and trifluoroacetic acid. The water used in this research was produced from a Millipore Milli-Q™ water system (Millipore Corporation, Milford, MA, USA). Ethanol (95% (v:v) and anhydrous) was obtained from Commercial Alcohols Inc. (Brampton, ON) through the College of Agriculture and Bioresources chemical stores (Saskatoon, SK).

### 2.3 Ethanol-formic acid-water phenolic extracts

Ethanol-formic acid-water extracts (EFW, 70:2:28% (v:v:v)) were prepared for each haskap variety using 25.00 ± 0.03 g of berry macerate. The fruit macerate was produced by mechanically blending (Osterizer™, Sunbeam Canada, Toronto, ON) at speed #10 for 2 min, followed by the addition of 50.00 ± 0.03 g EFW. The resulting mixture was covered and stirred at 700 rpm/4 ± 2°C for 2 h. The mixture was then vacuum filtered (#413, 12.5 cm; VWR International), and the remaining sediment was removed and resuspended in 50.00 ± 0.03 g EFW. This mixture was stirred at 4 ± 2°C for 2 h as outlined above and vacuum filtered a second time. The filtrates were combined and quantitatively transferred to a 200 mL volumetric flask and brought to volume with EFW. Extracts were stored in lightproof containers at −20 ± 2°C until analysis. Samples were extracted in triplicate.

### 2.4 Fractionation of EFW extracts

Amberlite^®^ XAD16N resin was hydrated in 50% aqueous ethanol (v:v) for 30 min then packed into a glass column (50 x 2.0 cm) to produce a resin bed of approximately 75 mL. The stationary phase was pre-conditioned with sequential treatment of 90 mL of water, 90 mL of 100% ethanol, and 90 mL of water at a flow rate of 2.5 mL/min. Individual 10.0 mL aliquots of haskap EFW extracts were rotary evaporated (35°C) (BUCHI Labortechnik AG, Switzerland), re-dissolved in 5.0 mL of water, and quantitatively transferred to the resin column. The stationary phase was then treated sequentially with 60 mL of water (fraction 1), 60 mL of 20% (v:v) ethanol (fraction 2), 60 mL of 40% (v:v) ethanol (fraction 3), 60 mL of 70% (v:v) ethanol (fraction 4), and 60 mL of 100% ethanol (fraction 5). Following fractionation, 3.0 mL of each fraction was individually removed and freeze dried (Heto-Holten A/S, Allerod, Denmark) then resuspended in 250 μL EFW prior to chromatographic analysis. The remaining volumes of each fraction were concentrated by rotary evaporation (35°C) and freeze dried. Freeze dried fractions were stored in lightproof containers at −20 ± °2C. Five fractionation experiments were conducted for each variety.

### 2.5 Phenolic rich extracts

Phenolic rich extracts were prepared by loading 10.0 mL of EFW extract (section 2.3) resuspended in 5.0 mL of water onto a prepared Amberlite^®^ XAD16N resin column (as outlined in section 2.4). The stationary phase was then treated with 135 mL of water to remove ascorbic acid, carbohydrates, and organic acids. Phenolics were then eluted with 135 mL of 100% ethanol. The resulting phenolic rich (PR) extracts were concentrated to dryness by rotary evaporation (35°C) and quantitatively transferred to a 10 mL volumetric flask and brought to volume with EFW. Sample extracts were transferred to lightproof containers and stored at −20 ± 2°C until analysis. All PR extracts were prepared in triplicate.

### 2.6 Total phenolic chromatographic index (TPCI)

Sample total phenolic chromatographic index (TPCI) was determined for EFW and PR extracts, and phenolic fractions employing high performance liquid chromatography with photodiode array detection (HPLC-PDA). Phenolics were identified and placed into specific phenolic subclasses (e.g. hydroxybenzoic acids, hydroxycinnamic acids, flavanols) based on UV-visible spectra comparison to reference standards. The concentration of each subclass was determined by chromatographic peak area comparison to reference standards and TPCI was determined by the summation of all subclass concentrations.

Chromatography was performed on an 1100 series HPLC system (Agilent Technologies Canada Incorporated, Mississauga, ON) with separation achieved using a 250 x 4.6 mm Prodigy ODS-3 5 μm, 100 Å, C_18_ endcapped column (Phenomenex) in series with a C_18_ guard cartridge (Phenomenex) at 25.0 ± 1.0 °C. The gradient mobile phase system used for phenolic compound separation consisted of 10 mM formic acid (solvent A) and 70:30 (v:v) acetonitrile:solvent A (solvent B). The linear gradient program was as follows: 100% A for 3 min, to 4% B at 6 min, to 10% B at 15 min, to 15% B at 30 min, to 20% B at 35 min, to 23% B at 50 min, to 25% B at 60 min, to 30% B at 66 min, to 50% B at 80 min, to 80% B at 85 min, which was held at 80% B for 5 min. For EFW and PR extracts, the injection volume was 20 μL; for fraction analysis, the injection volume was 50 μL. The mobile phase flow rate was 0.8 mL/min. All samples were syringe filtered (nylon, 0.2 μm pore size, 13 mm diameter, Chromatographic Specialties, Brockville, ON) prior to HPLC analysis. Phenolic compound detection was achieved employing a PDA detector with monitoring at 254, 280, 360, and 520 nm, with reference at 360, 400, 700, and 700 nm, respectively. Phenolic standards, apigenin, arbutin, caffeic acid, catechin, chlorogenic acid, cyanidin-3-*O*-glucoside, cyanidin-3-*O*-rutinoside, epicatechin, ferulic acid, 4-hydroxybenzoic acid, gallic acid, naringenin, *p*-coumaric acid, phloridzin, quercetin, rutin, and vanillic acid, were prepared at 100.0 ± 0.2 mg/L and were used to identify sample spectral (i.e. UV-visible profiles), retention time (RT), and quantitation parameters. The representative compounds selected for each subclass were: anthocyanins, cyanidin-3-*O-*glucoside; flavanols, catechin; flavanones, naringenin; flavonols, rutin; hydroxybenzoic acids, gallic acid; and hydroxycinnamic acids, chlorogenic acid. For quantification, standard curves were prepared at concentrations ranging from 0.5 to 2500 mg/L and had correlation coefficients ≥0.980. All samples were analyzed in triplicate.

### 2.7 Phenolic mass spectrometry

Supplementation of phenolic compound identification by HPLC-PDA was afforded by tandem MS employing an Agilent 1220 series HPLC coupled with an API QSTAR XL MS/MS hybrid QqToF tandem mass spectrometer (MS) equipped with an ESI source (Applied Biosystems Inc., CA, USA). The HPLC gradient system was the same as described previously (section 2.6). Nitrogen was used as both the drying and ESI nebulizing gas. External calibration employing caesium iodide (m/z 132.9054) and sex pheromone inhibitor iPD1 (m/z 829.5398) were used to ensure high mass accuracies. The ESI source was operated in the negative ion mode, and a sample injection volume of 20 μL was employed. MS/MS spectra were acquired using collision energies of −30, −45, and −55 v and analyzed using Analyst software (version 1.62).

### 2.8 Quantitative anthocyanin analysis

Sample EFW extracts were analyzed by high performance liquid chromatography with photodiode array detection (HPLC-PDA). Anthocyanin separation was afforded using a 250 x 4.6 mm Prodigy ODS-3 5 μm, 100 Å, C_18_ endcapped column (Phenomenex). The mobile phase system consisted of aqueous 4.0% (v:v) phosphoric acid at pH 1.4 (solvent A) and acetonitrile (solvent B) with the following gradient conditions: initial, 6% B for 12 min, followed by a linear gradient to 20% B at 66 min, and then held at 20% B for 18 min. The mobile phase flow rate was 0.8 mL/min. The sample injection volume was 20 μL, and all samples were syringe filtered prior to analysis. Analyte detection was achieved using a PDA detector with monitoring at 520 nm and reference at 700 nm. Anthocyanin standards used for identification were: cyanidin-3-*O*-arabinoside, cyanidin-3-*O*-galactoside (ideain), cyanidin-3-*O*-glucoside (kuromanin), cyanidin-3,5-*O*-diglucoside, cyanidin-3-*O*-rutinoside (keracyanin), cyanidin-3-*O*-xyloside, pelargonidin-3-*O*-glucoside, and peonidin-3-*O*-glucoside. Sample anthocyanin identification was afforded by retention time comparison to standards and spiking experiments, and standard curves (0.5-100 mg/L; correlation coefficients ≥0.980) were employed to determine their concentrations. All samples were analyzed in triplicate.

### 2.9 Anthocyanin mass spectrometry

Anthocyanin separation was afforded using the previously described stationary phase (section 2.8) with a mobile phase system (Palíková et al., 2008) consisting of aqueous 0.12% (v:v) trifluoroacetic acid with 5% (v:v) acetonitrile (solvent A) and 0.12% (v:v) trifluoroacetic acid in 100% acetonitrile (solvent B) with the following gradient conditions: initial, 5% B for 5 min, to 14% B at 50 min, to 20% at 55 min, to 5% B at 60 min, and held at 5% to 65 min. This separation method was used in tandem with the MS instrumentation described previously (section 2.7). Nitrogen was used as both the drying and ESI nebulizing gas. External calibration employing caesium iodide (m/z 132.9054) and sex pheromone inhibitor iPD1 (m/z 829.5398) were used to ensure high mass accuracies. The ESI source was operated in the positive ion mode with a sample injection volume of 20 μL. MS/MS spectra were acquired using collision energies of 20, 25, and 30 v and analyzed using Analyst software (version 1.62). Anthocyanins were also detected employing a photodiode array (PDA) detector with monitoring at 520 nm and reference at 700 nm. The Tundra variety 70% ethanol fraction was analyzed with a 20 μL injection volume.

### 2.10 ABTS (2,2’-azino-bis-3-ethylbenzthiazoline-6-sulfonic acid) radical scavenging assay

A radical cation (ABTS^•+^) solution was prepared by mixing 4.00 mL of 7.0 mM ABTS and 2.00 mL of 7.0 mM potassium persulfate in water. This mixture was maintained at room temperature for 12 h in the dark to afford radical ABTS^•+^ formation. The resulting ABTS^•+^ radical cation solution was then diluted using 70% methanol (v:v) to achieve a solution with an absorbance reading of 0.75 ± 0.05 at 734 nm (control). Trolox standards were prepared at concentrations of 0.1 mg/mL (0.4 mM) to 0.5 mg/mL (2.0 mM). Samples were diluted using 70% methanol to obtain absorbance values within the standard curve. EFW and PR extracts were diluted directly; 40, 70, and 100% ethanol fractions were prepared by combining the freeze dried material from 5 column fractionations (section 2.4) then re-dissolving in EFW prior to dilution with 70% methanol. The assay was conducted by individually mixing 10 μL of each sample dilution, control (70% methanol), standard (i.e. Trolox), with 1.00 mL of ABTS^•+^ solution. All solutions were vortexed and then stored in the dark for 6 min, followed by spectroscopic measurement at 734 nm. Percent ABTS radical scavenging activity was calculated as follows:

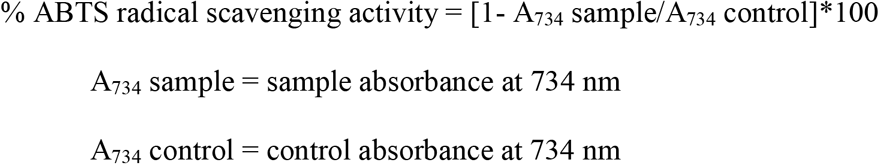

The % ABTS^•+^ inhibition was plotted as a function of sample concentration and linear regression equations were determined. Correlation coefficients of the linear regression equations were ≥0.950. All samples were analyzed in triplicate. The % ABTS^•+^ inhibition of 1.0 mM Trolox was determined from linear regression curves of the Trolox standards. The sample concentration equivalent to the inhibition activity of 1.0 mM Trolox was calculated. The Trolox equivalent antioxidant capacity (TEAC) was expressed as the equivalent activity of Trolox (mM)/100 mg sample as follows:

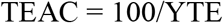

TEAC = Trolox equivalent antioxidant capacity (Trolox equivalents/100 mg sample); 100 = conversion factor to standardize the sample to 100 mg/mL; YTE = sample concentration (mg/mL) producing an ABTS^•+^ inhibition equivalent to 1 mM Trolox

### 2.11 DPPH (2,2-diphenyl-1-picrylhdrazyl) radical scavenging assay

A 500 μM DPPH solution was prepared in 70% methanol (v:v) with sonication for 60 min (4 ± 2 °C; Bransonic, Danbury, CT, USA). Fresh DPPH was prepared daily for sample analysis. Samples were diluted with 70% methanol prior to analysis to achieve DPPH radical scavenging of approximately 10 to 85%. A 250 μL aliquot of the diluted sample was added to 1.00 mL of DPPH solution. Samples were then vortexed and stored in the dark for 15 min before their absorbance at 517 nm was determined. A blank of 70% aqueous methanol was used. Percent DPPH radical scavenging activity was calculated as follows:

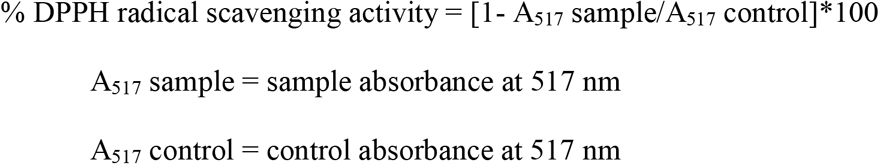

The 50% radical inhibition concentration (IC_50_) was determined by plotting the % DPPH radical scavenging versus concentration for each sample by linear regression. The IC_50_ value was expressed as mg solids/mL of DPPH solution and the antioxidant activity was reported as 1/ IC_50_. Regression equations had correlation coefficients ≥ 0.950. All samples were analyzed in triplicate.

### 2.12 Rancimat analysis

The ability of a phenolic rich extract (Tundra variety) to delay oxidation in a polyunsaturated oil (borage) was determined employing a 697 Rancimat (AOCS Official Method Cd 12b-92). Samples analyzed included, borage oil, borage oil + 0.01 and 0.02% BHT (w:w), borage oil + 0.1 and 0.2% Rosamox (w:w), and borage oil + 500 ppm Tundra phenolic rich extract. All samples were sonicated for 30 min prior to Rancimat analysis to completely disperse the antioxidants. Samples (3.00 ± 0.02 g) were heated to 110°C and dry air at a flow rate of 20 L/h was passed through the mixture.

### 2.13 Sample preparation and analysis flow chart

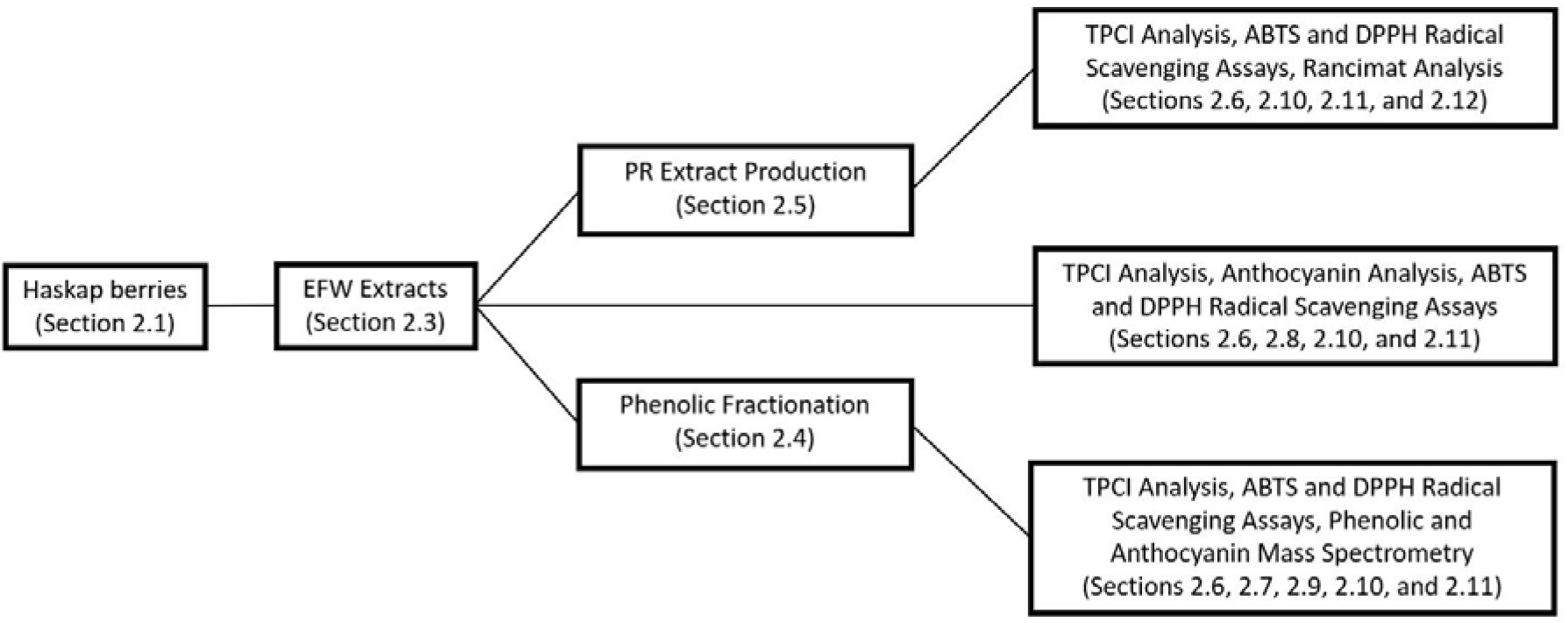

### 2.14 Statistical analysis

Statistical analysis of chemical experimental data was performed using SPSS software for Windows version 22.0 (IBM SPSS Inc., Chicago, IL, USA) for one-way analysis of variance (ANOVA). Difference between means (p<0.05) was determined using the multiple-comparison Tukey’s HSD (honestly significant difference) multiple comparison test.

## 3. Results & discussion

Phenolic concentration, subclass composition, and chromatographic separation were determined for the Aurora, Blizzard, Honey Bee, Indigo Gem, and Tundra haskap varieties, which were bred and grown in Saskatoon, Saskatchewan (Canada). Structural analysis by mass spectrometry was also performed on the Tundra variety. Solid phase extraction (SPE) was employed to create a phenolic rich (PR) extract (>99% phenolics) and a series of resin-ethanol fractions, which contained haskap phenolics separated based on structure (i.e. subclass). Haskap phenolic extracts and fractions were subjected to *in vitro* free radical scavenging (RS) assays and rancimat analysis to determine their antioxidant activities and potential to delay lipid oxidation for industrial and health purposes.

### 3.1 Total phenolic chromatographic indices (TPCI) of haskap phenolic extracts

The total phenolic chromatographic index (TPCI) of a sample is defined as the sum of all extracted phenolics as analyzed by HPLC-PDA and is calculated by summation of all identified phenolic subclasses found in the sample (Escarpa & Gonzalez, 2001). Initially, the phenolic subclass for each peak was identified by UV-visible spectrum comparison to standards, followed by subclass concentration determination using peak area summation and comparison with a representative phenolic of that subclass through linear regression (i.e. concentration *vs*. peak area). Reference standards included catechin, chlorogenic acid, cyanidin-3-*O*-glucoside, gallic acid, naringenin, and rutin, which were representatives of the flavanol, hydroxycinnamic acid, anthocyanin, hydroxybenzoic acid, flavanone, and flavonol subclasses, respectively.

Sample compounds with a detector response at 280 nm (i.e. peaks with a signal to noise ratio ≥3x) and a UV-visible spectrum matching those of phenolic standards, were assumed to be phenolics and included in the TPCI analysis. The anthocyanin and flavonol subclasses were quantified at wavelengths of 520 and 360 nm, respectively, with all other phenolic subclasses at 280 nm. The TPCI values for the ethanol-formic acid-water (EFW) extracts of each haskap berry variety are shown in Table 1, and a representative HPLC-PDA chromatogram of the EFW phenolic extract for the Tundra variety is shown in Fig. 1.

**Table 1.** Mean and standard deviation phenolic subclass and total TPCI results for the ethanol:formic acid: water (EFW) extracts of Aurora, Blizzard, Honey Bee, Indigo Gem, and Tundra haskap berry varieties.

|  | Aurora | Blizzard | Honey Bee | Indigo Gem | Tundra |
| --- | --- | --- | --- | --- | --- |
| Hydroxybenzoic acids | 7.5 ± 0.3<sup>2,3</sup> | 14.9 ± 0.7 | 19.3 ± 2.2 | 4.7 ± 2.2 | 9.8 ± 0.2 |
| Flavanols | 71.6 ± 2.2 | 23.6 ± 0.7 | 143.1 ± 1.9 | 49.5 ± 1.2 | 130.9 ± 4.0 |
| Hydroxycinnamic acids | 46.5 ± 1.7 | 58.2 ± 1.7 | 56.4 ± 1.5 | 56.6 ± 1.6 | 80.2 ± 1.2 |
| Flavonols | 60.3 ± 6.3 | 29.8 ± 1.1 | 38.1 ± 1.2 | 90.7 ± 2.7 | 68.9 ± 1.6 |
| Flavanones | 7.4 ± 0.3 | 10.6 ± 0.5 | 0.6 ± 0.1 | 9.9 ± 0.2 | 7.8 ± 0.1 |
| Anthocyanins | 249.2 ± 8.3 | 315.9 ± 1.8 | 253.5 ± 4.4 | 447.8 ± 5.9 | 429.4 ± 6.3 |
| TPCI<sup>1</sup> | 442.5 ± 14.5 | 453.1 ± 3.0 | 511.2 ± 3.3 | 659.2 ± 10.5 | 727.0 ± 3.6 |

<sup>1</sup>Total Phenolic Chromatographic Index = sum of all identified and quantified phenolic peaks.
<sup>2</sup>mg/100 g FW.
<sup>3</sup>Mean ± standard deviation results of triplicate sample analysis.

**Fig. 1.**
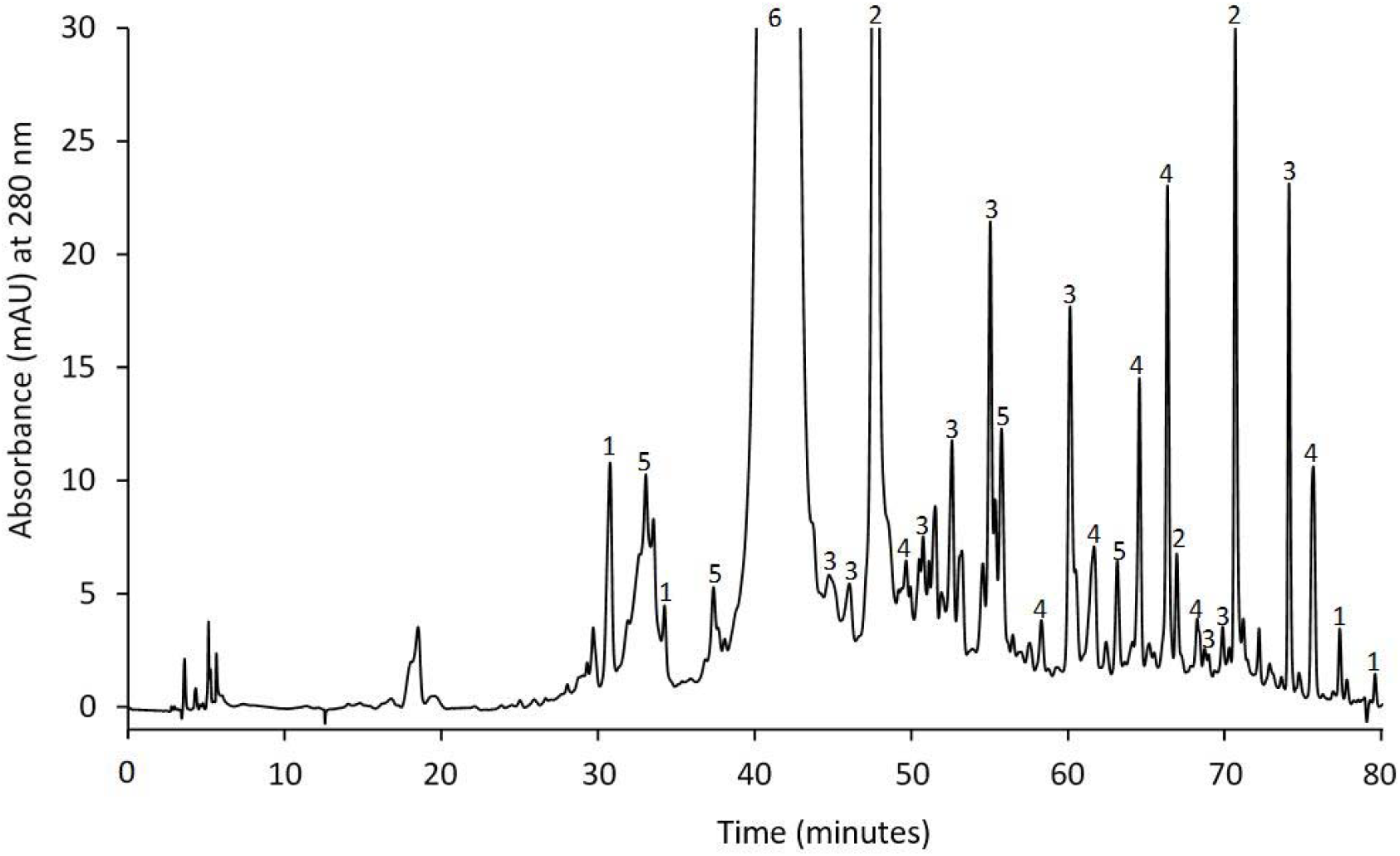
Representative HPLC-PDA chromatogram of the Tundra variety showing the identification of peak phenolic subclasses in an EFW extract. Peak phenolic subclass assignments: 1. hydroxybenzoic acids; 2. hydroxycinnamic acids; 3. flavanols; 4. flavonols; 5. flavanones; 6. anthocyanins.

The TPCI values for the EFW extracts of the five haskap samples ranged from 442.5-727.0 mg/100 g FW. Tundra variety haskaps were found to have the highest phenolic content by TPCI and Aurora the lowest. Variation in fruit phenolic content can be expected due to differences in climate (temperature, rainfall, sunlight) and agricultural practices (fertilization, watering practices); however, for these five varieties grown in the same location and under the same growing practices, these results highlight variation primarily due to genetic differences. These results are consistent with other published works which have reported high phenolic content in Indigo Gem (8.42 ± 0.02 mg GAE/g FW) and Tundra (8.09 ± 0.03 mg GAE/g FW) varieties using a total phenolic content (TPC) assay (Khattab, Ghanem, & Brooks, 2016).

An important physical property of berries/fruits that have a significant impact on their total phenolic content is size (i.e. surface area:volume ratio) as phenolics, particularly pigments such as anthocyanins, are generally localized in the skin or peel (Bakowska-Barczak, Marianchuk, & Kolodziejckz, 2007; Ribeiro de Souza, Willems, & Low, 2019). The EFW extract TPCI results for these five varieties were directly related to berry size with Tundra (17.41 ± 1.65 mm) being the smallest of the five varieties and having the highest TPCI value (727.0 ± 3.6 mg/100 g FW). Concomitantly, the Aurora variety (23.49 ± 3.23 mm) had the largest size and the lowest TPCI value (442.5 ± 14.5 mg/100 g FW).

The major phenolic subclass identified in the haskap EFW extracts by TPCI analysis was the anthocyanins which accounted for 49.6-69.7% of the TPCI values. Indigo Gem and Tundra had the highest anthocyanin contents of 447.8 ± 5.9 and 429.4 ± 6.3 mg/100 g FW mg/100 g FW, respectively. Because the anthocyanin subclass was responsible for the largest proportion of haskap TPCI, anthocyanin and TPCI results were strongly related. In addition to anthocyanins, flavanols (5.2-28.0%), hydroxycinnamic acids (8.6-12.8%), and flavonols (6.6-13.8%) were identified as the next most abundant phenolic subclasses by TPCI with lower levels of hydroxybenzoic acids (0.7-3.3%) and flavanones (0.1-2.3%) observed. These results align with literature which has reported major phenolic classes in haskap berries to include flavonoids (e.g. flavanols and flavonols) and phenolic acids (e.g. hydroxycinnamic and hydroxybenzoic acids) (Palíková et al., 2008; Jurikova et al., 2012; Wojdyło, Jáuregui, Carbonell-Barrachina, Oszmiański, & Golis, 2013; Khattab, Brooks, & Ghanem, 2015; Rupasinghe et al., 2015; Kucharska, Sokół-Łętowska, Oszmiański, Piórecki, & Fecka, 2017; Senica, Bavec, Stampar, & Mikulic-Petkovsek, 2018).

### 3.1 Total qualitative and quantitative anthocyanin analysis by HPLC-PDA

Qualitative and quantitative analysis of EFW extracts by HPLC-PDA identified the major anthocyanin as cyanidin-3-*O-*glucoside (retention time [RT]: 46.0 min) which accounted for 73.1-82.5% of the total anthocyanins. Other identified anthocyanins included cyanidin-3,5-*O-* diglucoside (RT: 38.5 min), cyanidin-3-*O-*rutinoside (RT: 48.2 min), pelargonidin-3-*O-*glucoside (RT: 50.1 min), peonidin-3-*O-*glucoside (RT: 53.9 min), and cyanidin-3-*O-*xyloside (RT: 56.2 min) (Table 2). A representative HPLC-PDA chromatogram can be found in Supplementary material Fig. S1A.

**Table 2.** Mean and standard deviation anthocyanin contents for Aurora, Blizzard, Honey Bee, Indigo Gem, and Tundra haskap berry varieties as determined by HPLC-PDA.

|  | Aurora | Blizzard | Honey Bee | Indigo Gem | Tundra |
| --- | --- | --- | --- | --- | --- |
| Cyanidin-3,5-*O*-diglucoside | 17.5 ± 0.3<sup>1,2a</sup> | 34.2 ± 0.5<sup>c</sup> | 48.1 ± 1.4<sup>d</sup> | 47.3 ± 1.8<sup>d</sup> | 30.3 ± 1.6<sup>b</sup> |
| Cyanidin-3-*O*-galactoside | ND<sup>3</sup> | ND | ND | ND | ND |
| Cyanidin-3-*O*-glucoside | 227.8 ± 4.8<sup>a</sup> | 269.4 ± 1.3<sup>b</sup> | 227.3 ± 1.9<sup>a</sup> | 396.3 ± 12.4<sup>d</sup> | 361.6 ± 11.9<sup>c</sup> |
| Cyanidin-3-*O*-rutinoside | 13.2 ± 0.4<sup>a</sup> | 28.6 ± 0.9<sup>c</sup> | 22.7 ± <0.1<sup>b</sup> | 32.4 ± 0.38<sup>d</sup> | 26.3 ± 1.7<sup>c</sup> |
| Pelargonidin-3-*O*-glucoside | 2.3 ± 0.1<sup>b</sup> | 1.1 ± <0.1<sup>a</sup> | 1.1 ± 0.1<sup>a</sup> | 4.9 ± 0.2<sup>c</sup> | 1.0 ± 0.1<sup>a</sup> |
| Peonidin-3-*O*-glucoside | 15.4 ± 0.1<sup>b</sup> | 20.9 ± 0.8<sup>c</sup> | 11.8 ± 0.2<sup>a</sup> | 20.4 ± 0.3<sup>c</sup> | 31.8 ± 0.9<sup>d</sup> |
| Cyanidin-3-*O*-xyloside | NR<sup>4</sup> | NR | NR | NR | NR |
| Total | 276.2 ± 5.0<sup>b</sup> | 354.3 ± 2.7<sup>c</sup> | 311.0 ± 1.0<sup>b</sup> | 501.3 ± 13.5<sup>e</sup> | 451.0 ± 13.5<sup>d</sup> |

<sup>1</sup>mg/100 g FW.
<sup>2</sup>Mean ± standard deviation results of triplicate sample analysis.
<sup>3</sup>Not detected (<0.8 mg/100 g FW).
<sup>4</sup>Not reported due to coelution.
<sup>a-c</sup>Mean values with a common letter were not statistically different (p<0.05) by Tukey's HSD multiple range test between berry varieties.

The Indigo Gem variety contained the highest total anthocyanin content (501.3 ± 13.5 mg/100 g FW) along with the highest concentration of cyanidin-3-*O-*glucoside (396.3 ± 12.4 mg/100 g FW). The Tundra variety also had high anthocyanin (451.0 ± 13.5 mg/100 g FW) and cyanidin-3-*O-*glucoside concentrations (361.6 ± 11.9 mg/100 g FW). The anthocyanin composition and range for the five varieties were: 6.3-15.5% cyanidin-3,5-*O-*diglucoside, 73.1-82.5% cyanidin-3-*O-*glucoside, 4.8-8.1% cyanidin-3-*O-*rutinoside, 3.8-7.1% peonidin-3-*O-* glucoside, and 0.2-1.0% pelargonidin-3-*O-*glucoside. In addition, cyanidin-3-*O-*xyloside was identified qualitatively in all varieties, but was not quantified either due to low concentrations (i.e. <6x signal to noise ratio) and/or coelution with other anthocyanins/compounds (i.e. absorbance at 520 nm).

These results agreed with those reported by Khattab et al., (2016) where anthocyanins were identified as the major phenolic subclass, with total concentrations of 6.40 ± 0.13 mg/g FW for Tundra and 7.09 ± 0.24 mg/g FW for Indigo Gem varieties based on HPLD-PDA quantification. However, literature results for Indigo Gem and Tundra varieties employing the pH-differential method were much lower than those reported in this work at 246.3 ± 2.8 and 303.2 ± 5.5 mg CGE/100 g FW, respectively (Rupasinghe et al., 2015). These authors also reported much lower anthocyanin contents for Indigo Gem and Tundra varieties of 178.9 and 214.5 mg/100 g FW, respectively by HPLC-PDA.

### 3.2 Antioxidant activities of haskap berry extracts

Haskap berry EFW extracts (section 2.3) and solid phase extraction fractions isolated from the Tundra variety (section 2.4) were examined for their free radical scavenging (RS) abilities employing two *in vitro* assays: ABTS and DPPH. The radical scavenging (RS) EFW extract results for all varieties and the Tundra variety resin-ethanol fractions are reported in Table 3. The highest EFW phenolic extract RS value for ABTS was found for the Tundra variety of 225.9 ± 5.8 mM Trolox equivalents (TEAC)/100 mg FW, which was significantly higher than those observed for the other four varieties (range of 131.4-207.0). In addition, the 1/IC_50_ DPPH value for the Tundra variety of 32.7 ± 1.3 was significantly higher than all but the Indigo Gem variety (31.3 ± 0.8). These *in vitro* RS results align with the Tundra and Indigo Gem varieties having the highest TPCI and total anthocyanin values (Tables 1 and 2). Tundra solid phase extraction fraction RS results are discussed in Section 3.3.

**Table 3.** Mean and standard deviation ABTS and DPPH radical scavenging activity data for EFW extracts of Aurora, Blizzard, Honey Bee, Indigo Gem, and Tundra haskap berry varieties and the solid phase chromatographic fractions from the Tundra variety.

|  | ABTS Radical Scavenging<br>Activity<sup>1</sup> | DPPH Radical Scavenging<br>Activity<sup>2</sup> |
| --- | --- | --- |
| Aurora | 131.4 ± 5.9<sup>3a</sup> | 23.7 ± 1.0<sup>a</sup> |
| Blizzard | 158.5 ± 2.5<sup>b</sup> | 24.8 ± 0.7<sup>a</sup> |
| Honey Bee | 146.3 ± 3.0<sup>b</sup> | 23.6 ± 0.4<sup>a</sup> |
| Indigo Gem | 207.0 ± 5.2<sup>c</sup> | 31.3 ± 0.8<sup>b</sup> |
| Tundra | 225.9 ± 5.8<sup>d</sup> | 32.7 ± 1.3<sup>b</sup> |
| 40% ethanol fraction<sup>4</sup> | 56.2 ± 1.5 | 9.4 ± 0.6 |
| 70% ethanol fraction<sup>4</sup> | 145.0 ± 4.6 | 14.8 ± 1.3 |
| 100% ethanol fraction<sup>4</sup> | 6.4 ± 0.1 | 1.3 ± <0.1 |

<sup>1</sup>Reported as mM Trolox activity equivalent (TEAC)/100 mg FW.
<sup>2</sup>Reported as 1/IC<sub>50</sub> (1/100 mg FW for 50% DPPH radical inhibition).
<sup>3</sup>Mean ± standard deviation results of triplicate sample analysis.
<sup>4</sup>Tundra variety.
<sup>a-c</sup>Mean values in the same column followed by a common letter were not statistically different (p<0.05) by Tukey's HSD multiple range test between the varieties.

When compared to literature, the *in vitro* RS results from this study were higher than: (a) the ABTS value of 9.55 TEAC/100 g FW for an unidentified haskap variety grown in Alberta, Canada (Bakowska-Barczak et al., 2007); (b) the reported range of 12.65-49.73 TEAC/100 g dry matter (DM) for eight Polish varieties (Wojdyło et al., 2013); and (c) the DPPH results for Borealis, Indigo Gem, and Tundra varieties with 1/IC_50_ values ranging from 12.8-17.2 with a mean of 15.2 (Rupasinghe, Yu, Bhullar, & Bors, 2012). Variation in RS abilities can be impacted by analytical methods (i.e. reaction time and temperature), phenolic extraction technique (e.g. solvent), and sample phenolic content, which is dependent on variety, environmental growth conditions, and geographical origin. A number of literature citings of *in vitro* RS values for haskaps report their results in gallic acid equivalents (GAE) or % scavenging of the radical (i.e. for the DPPH assay) making direct comparison of values invalid (Kusznierewicz et al., 2012; Khattab et al., 2016; Kucharska et al., 2017).

### 3.3 Phenolic Analysis of Tundra haskap berry resin-ethanol fractions

Tundra haskap EFW extracts were fractionated using Amberlite^®^ XAD16N resin with increasing concentrations of aqueous ethanol for phenolic elution (i.e. 20, 40, 70, and 100% ethanol). The goal was to produce fractions containing higher proportions and/or separation of phenolic subclass(es), as determined by HPLC-PDA and HPLC-MS/MS. Chromatographic (HPLC-PDA) results showed that the water (100%) and 20% ethanol fractions did not contain significant phenolic concentrations (<5.6 mg/100 g FW; data not shown) and were not analyzed further. Representative chromatograms of the Tundra phenolic resin-ethanol fractions (40, 70, and 100% ethanol) are shown in Supplementary material, Fig. S1B.

Based on the total combined TPCI of all fractions (data not shown), the 40% ethanol fraction contained 27.4% of sample phenolics, which were primarily the anthocyanin and flavanol subclasses. The 70% ethanol fraction contained 63.8% of the phenolics, primarily anthocyanins along with flavonols, hydroxycinnamic acids, and flavanols. The 100% ethanol fraction contained 6.8% of the phenolics with flavanols and flavonols as primary components and a very low concentration of anthocyanins. The RS abilities of the fractions aligned with their phenolic content (Tables 1 and 2) with the 70% ethanol fraction being the most successful radical scavenger in both the ABTS and DPPH assays (Table 3). The 40% ethanol fraction was the next best radical scavenger with significantly lower values than the 70% fraction, and the 100% fraction had very low RS abilities for both assays. The values for these *in vitro* assays are reported in terms of fresh weight (FW). Therefore, the low RS ability of the 100% fraction is more indicative of the low overall concentration of these phenolics in the whole berry rather than the RS ability of the specific compounds present.

Results obtained from HPLC-MS/MS analysis of the 40, 70, and 100% ethanol fractions are shown in Table 4 with the corresponding labeled HPLC-PDA chromatograms in Supplementary material, Fig. S1B. With a combination of HPLC-PDA and HPLC-MS/MS analytical techniques, 22 non-anthocyanin phenolics were identified in the Tundra variety haskaps, primarily of the flavonol and hydroxycinammic acid subclasses. In some cases, glycosylated phenolics were identified that could not be fully characterized due to the potential for multiple isomers to produce the same fragmentation pattern. As an example, at 69.81 min in the 100% fraction (Table 4), a glycosylated kaempferol derivative was identified based on the precursor *m/z* 447 along with the characteristic product ions at *m/z* 284, 255, and 151. This fragmentation pattern could be produced by a molecule with a glucoside, galactoside, or other 6-carbon glycosylation. Without an available standard of each, the exact identity could not be confirmed and the phenolic was therefore identified more generically as a kaempferol-hexoside. Additionally, a selection of phenolics were identified that were not members of the six subclasses used for TPCI classification. These compounds were grouped together and labeled as other flavonoids/7 in Supplementary material, Fig. S1B and Table 4.

**Table 4.**
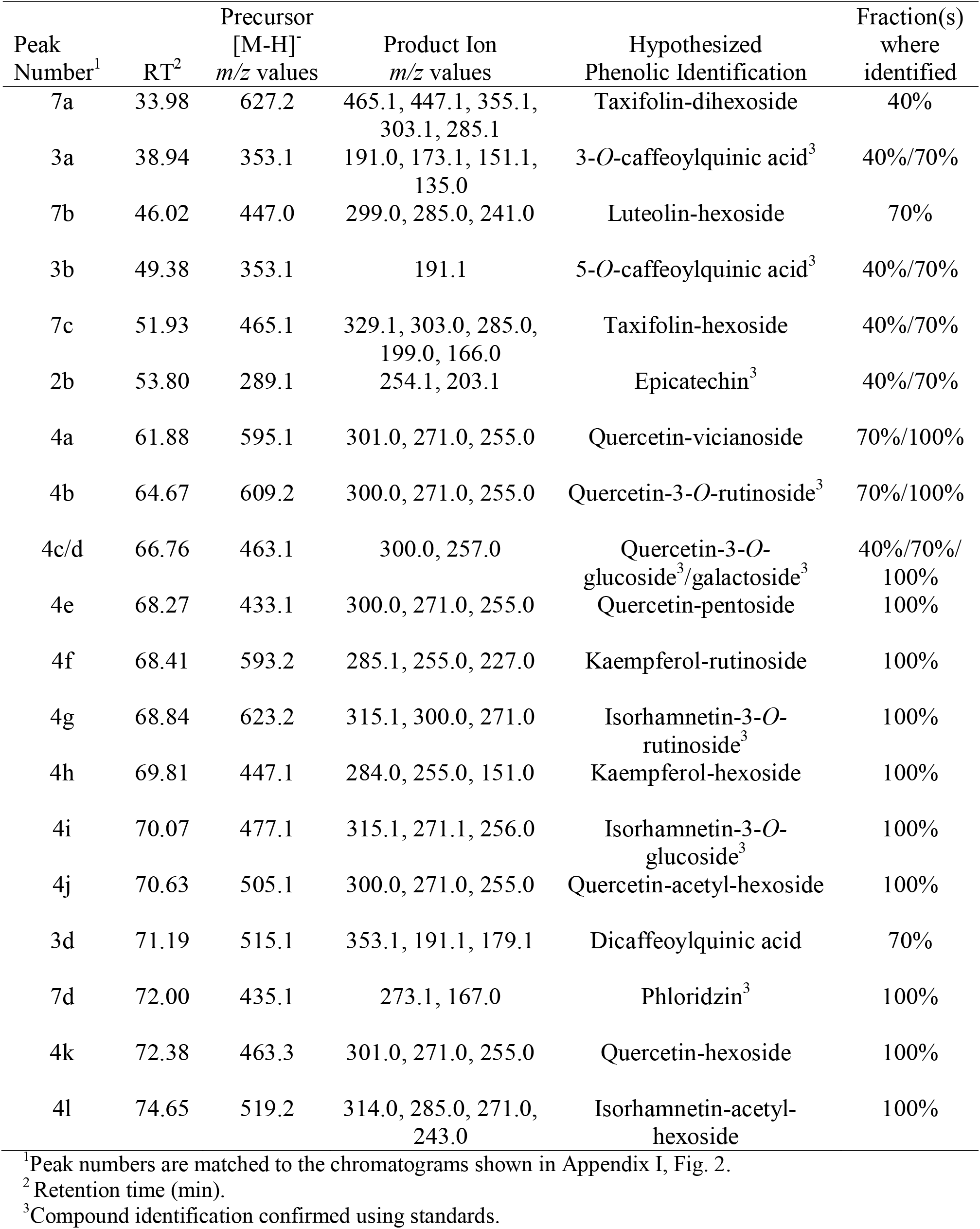
HPLC-MS/MS ion precursor and product ion *m/z* values and phenolic identification for the Tundra variety 40, 70, and 100% ethanol fractions.

Four phenolic acids were identified as: (a) 3-*O*-caffeoylquinic acid (neochlorogenic acid; 38.94 min); (b) 5-*O*-caffeoylquinic acid (chlorogenic acid; 49.38 min); (c) ferulic acid (62.15 min); and (d) a dicaffeoylquinic acid (71.19 min). The most abundant identified phenolic acid was 5-*O*-caffeoylquinic acid, which agrees with literature reports of the 3- and 5-*O-* caffeoylquinic acids being the most abundant in a selection of Russian and American (Oregon, USA) varieties (Chaovanalikit et al., 2004; Caprioli et al., 2016; Khattab et al., 2016).

Two flavanols, catechin and epicatechin, were identified, and these compounds have previously been reported as the most abundant flavanols in many Canadian-grown haskap varieties (Rupasinghe et al., 2015). For the flavonol subclass, the majority of those identified were quercetin derivatives, which included quercetin-3-*O*-galactoside, quercetin-3-*O*-glucoside, and quercetin-3-*O*-rutinoside, all of which have been reported in haskaps (Chaovanalikit et al., 2004; Kusznierewicz et al., 2012). In addition to these, literature has reported haskap varieties to contain many other kaempferol, isorhamnetin, and quercetin derivatives including those reported in Table 4 (Wojdyło et al., 2013; Khattab et al., 2015; Rupasinghe et al., 2015; Kucharska et al., 2017; Senica et al., 2018; Gołba, Sokół-Łętowksa, & Kucharska, 2020). Other flavonoids identified in the Tundra variety included taxifolin and luteolin glycosides along with phloridzen, all of which have been reported to be present in haskaps (Ochmian, Skupień, Grajkowski, Smolik, & Ostrowska, 2012; Kucharska et al., 2017).

### 3.4 Anthocyanin analysis of Tundra haskap berry 70% ethanol fraction

Due to the high concentration of anthocyanins in the haskap phenolic extracts, their identification by HPLC-MS/MS was conducted using a modified LC method (section 2.9) to afford compatibility with the mass spectrometer (i.e. trifluoracetic acid replacement of phosphoric acid). Anthocyanin results are reported in Table 5 with a corresponding HPLC-PDA chromatogram of the identified compounds in Supplementary material, Fig. S1C. This analysis confirmed the identification of the six anthocyanins previously reported in this work (Table 2), with agreement with the major anthocyanins found in haskaps as: cyanidin-3,5-*O*-diglucoside, cyanidin-3-*O-*glucoside, cyanidin-3-*O*-rutinoside, peonidin-3-*O*-glucoside, pelargonidin-3-*O*-glucoside, and cyanidin-3-*O*-xyloside (Chaovanalikit et al., 2004; Palíková et al., 2008; Kusznierewicz et al., 2012; Wojdyło et al., 2013; Khattab et al., 2015; Caprioli et al., 2016). Cyanidin-3-*O-*galactoside was not detected during the initial HPLC-PDA analysis but was seen with enhanced separation and detection via HPLC-MS/MS (Supplementary material, Fig. S1C).

**Table 5.**
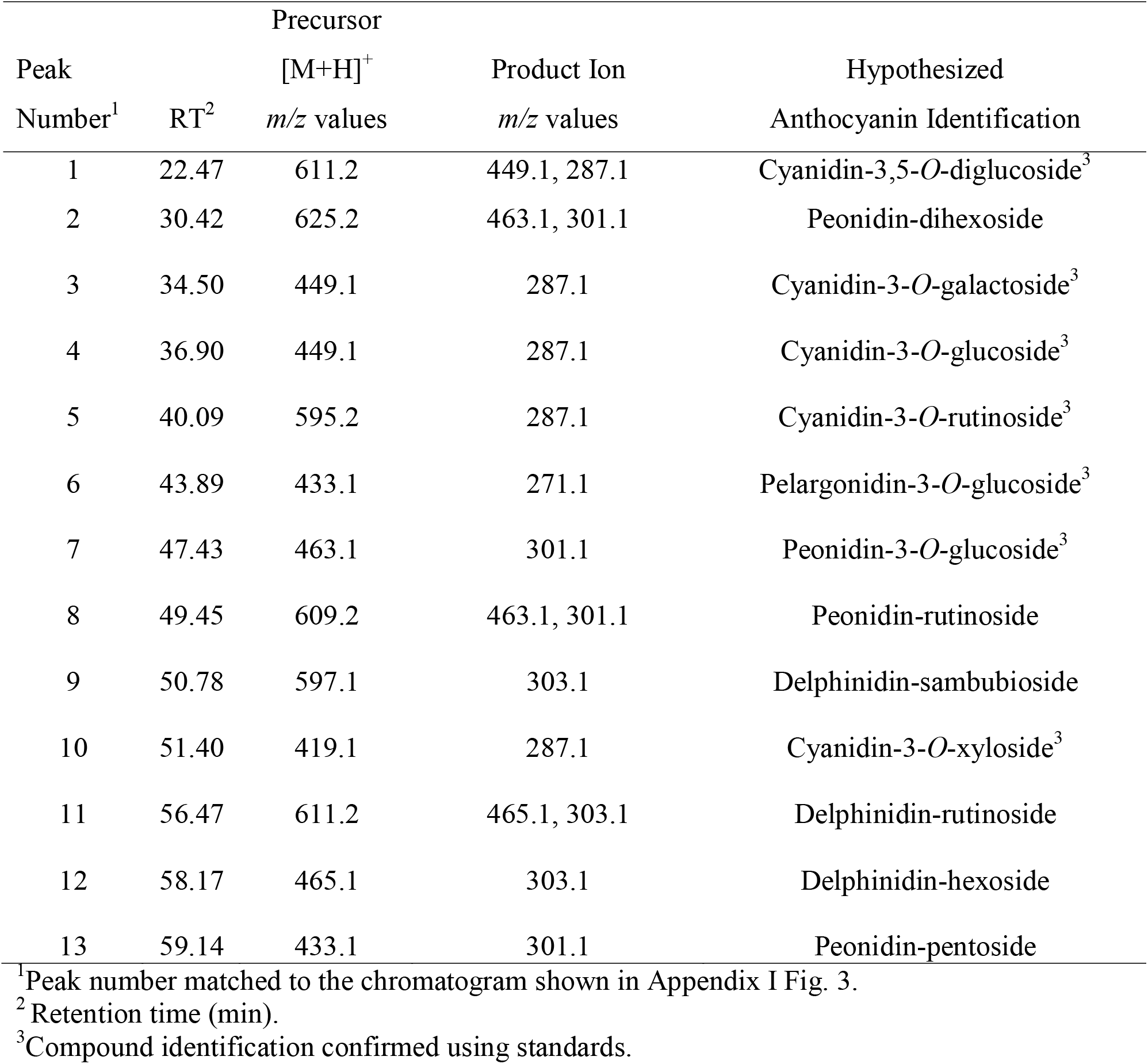
HPLC-MS/MS ion *m/z* values and compound identification obtained for anthocyanins in the Tundra variety 70% ethanol fraction.

A number of less common anthocyanins were identified in the Tundra variety studied via HPLC-MS/MS as: cyanidin-3-*O*-galactoside, cyanidin-3-*O*-xyloside, peonidin-dihexoside, peonidin-rutinoside, delphinidin*-*hexoside, and delphinidin-rutinoside. Cyanidin-3-*O*-galactoside and cyanidin-3-*O*-xyloside have been identified in haskaps grown in Alberta, Canada, of an unknown variety (Bakowska-Barczak et al., 2007). Delphinidin-derived anthocyanins are not often found in haskaps but have been reported in Czech Republic cultivars and more rarely in Canadian varieties including Tundra and Indigo Gem (Palíková et al., 2008; Rupasinghe et al., 2015). Two additional anthocyanins were tentatively identified by HPLC-MS/MS as delphinidin-sambubioside (β-D-xylosyl-(1→2)-β-D-glucose) and a peonidin-pentoside; neither compound has previously been reported in haskaps. Delphinidin-sambubioside was identified by the precursor *m/z* of 597 with the characteristic product of *m/z* 303 for the delphinidin aglycone; peonidin-pentoside was identified based on the peonidin product at *m/z* 301 and the precursor at *m/z* 433 which aligns with a mass difference for a 5-carbon carbohydrate (e.g. arabinose, xylose). Additional anthocyanins that have been reported in literature for haskaps include pelargonidin-3,5-*O-*diglucoside and pelargonidin-3-*O-*rutinoside which were not identified in this analysis of the Tundra variety (Palíková et al., 2008).

### 3.5 Rancimat analysis of Tundra haskap berry phenolics in borage oil

Rancimat analysis is a common technique used in the food industry to evaluate the oxidative stability of edible oils, and the potential for chemical compounds/mixtures to extend stability. Samples are heated (110 °C) under a stream of dry air and the rate of autoxidation is measured via changes in conductivity due to the formation of low molecular weight organic acids (Anotolovich et al., 2001). During this reaction, there is a sharp increase in the conductivity of the oxidation curve and a point of inflection is reached. A longer induction time represents improved stability of the oil/chemical compounds/mixtures. Rancimat analysis was conducted to determine the antioxidant abilities of the Tundra haskap PR extract (500 ppm) with comparison to the commercial antioxidants BHT (0.01 and 0.02%; w:w) and Rosamox (0.1% and 0.2%; w:w). Borage oil was selected as the reaction matrix as it is rich in polyunsaturated fatty acids (PUFAs) making it highly susceptible to oxidation (Khan & Shahidi, 2000).

The borage oil control had an induction time of 1.46 h, with results for the commercial antioxidant Rosamox at 0.1% (recommended minimum by the producer) and 0.2% showing minimal change in induction times of 1.38 and 1.44 h, respectively. The commercial antioxidant BHT at 0.01 and 0.02% lengthened induction times to 1.72 h and 2.18 h, respectively. The Tundra PR extract at 500 ppm was found to have the greatest effect on delaying oxidation with an induction time of 2.46 h, a 68% delay in the oxidation of borage oil. These results showed that a phenolic extract produced from haskap berries can be an effective natural antioxidant in edible oils rich in PUFAs.

## 4. Conclusions

The phenolic composition of five Saskatoon, Saskatchewan, bred and grown haskap berry varieties was determined using HPLC-PDA (TPCI and anthocyanins) and HPLC-MS/MS (for individual phenolic and anthocyanin identification). The most abundant phenolic subclasses in haskaps included anthocyanins (276.2-451.0 mg/100 g FW), primarily cyanidin-3-*O-*glucoside (227.3-396.3 mg/100 g FW), along with flavanols and flavonols. The Tundra variety had the highest phenolic content by TPCI, while Indigo Gem had the highest anthocyanin content. Two previously unreported anthocyanins were identified in the Tundra variety using HPLC-MS/MS as delphinidin-sambubioside and a peonidin-pentoside. Haskap phenolic extracts were also successfully fractionated using solid phase extraction in conjunction with an aqueous ethanol gradient to produce an anthocyanin rich fraction (40% ethanol) and a flavanol/flavonol rich fraction (100% ethanol). This allows for the potential isolation and concentration of the most active components in haskaps for nutraceutical use. Free radical scavenging assays (ABTS and DPPH) showed high radical scavenging abilities for each of the haskap extracts/fractions produced, and rancimat analysis of the Tundra phenolic rich extract significantly delayed lipid oxidation when compared to a commercial natural (Rosamox) and a synthetic antioxidant (BHT). Based on their high phenolic content and free radical scavenging ability, the phenolic extracts and fractions from haskap berries show significant potential as ingredients in food formulations, as natural antioxidants in polyunsaturated edible oils/oil-rich foods, and as nutraceutical products.

## Acknowledgements

This work was financially supported by the Saskatchewan Ministry of Agriculture (Agriculture Development Fund Grant 20190076); University of Saskatchewan (UofS) College of Graduate and Postdoctoral Studies and Department of Food and Bioproduct Sciences (LRZ), and the Natural Sciences and Engineering Research Council (NSERC) of Canada Discovery Grant Program (NHL: #36675; CHE: RGPIN 04930-2015). Fruit samples were provided by the UofS Horticulture Field Lab with the help of Dr. Bob Bors.

