## Supplemental Figures for "Phenolic profiles and antioxidant activities of Saskatchewan (Canada) bred haskap (*Lonicera caerulea)* berries"

**Supplementary material**

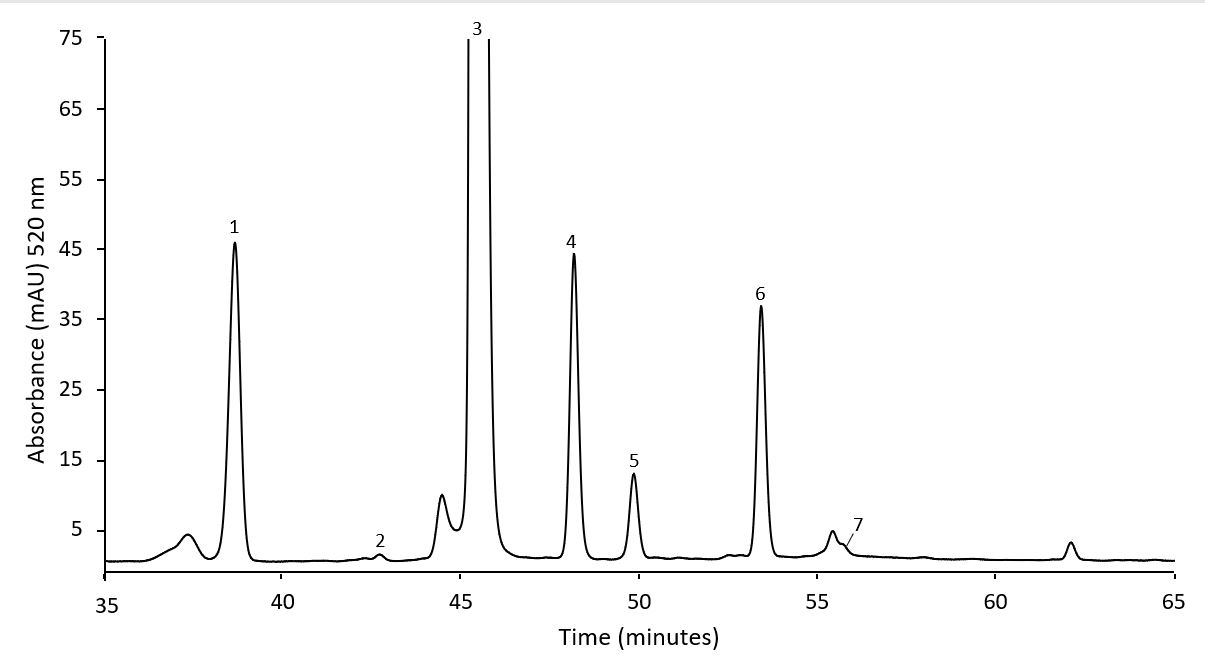

**Fig. S1A.** Representative HPLC-PDA chromatogram of the Indigo Gem variety showing the identification of anthocyanins. Peak assignments: 1. cyanidin-3,5-*O-*diglucoside; 2. cyanidin-3-*O-*glucoside; 3. cyanidin-3-*O-*rutinoside; 4. pelargonidin-3-*O-*glucoside; 5. peonidin-3-*O-*glucoside; and 6. cyanidin-3-*O-*xyloside.

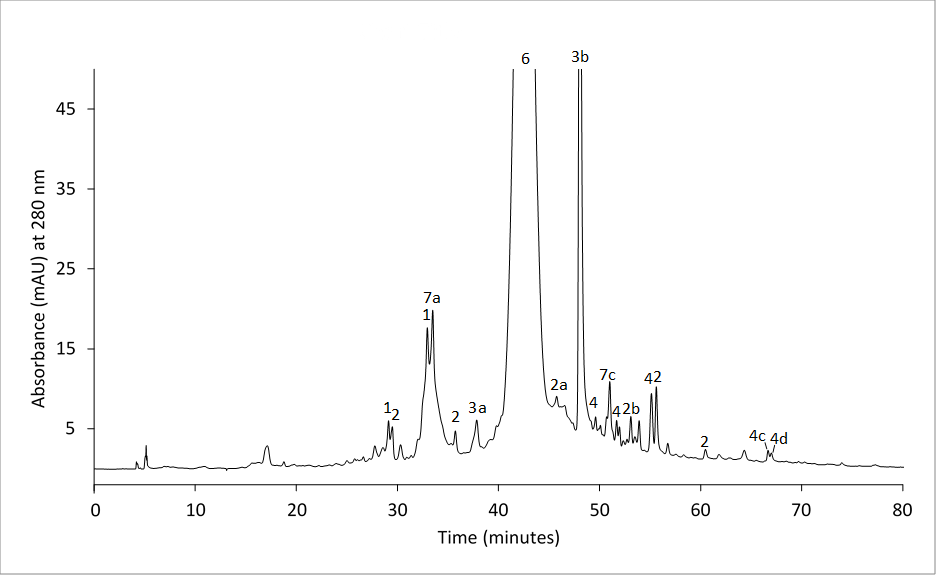

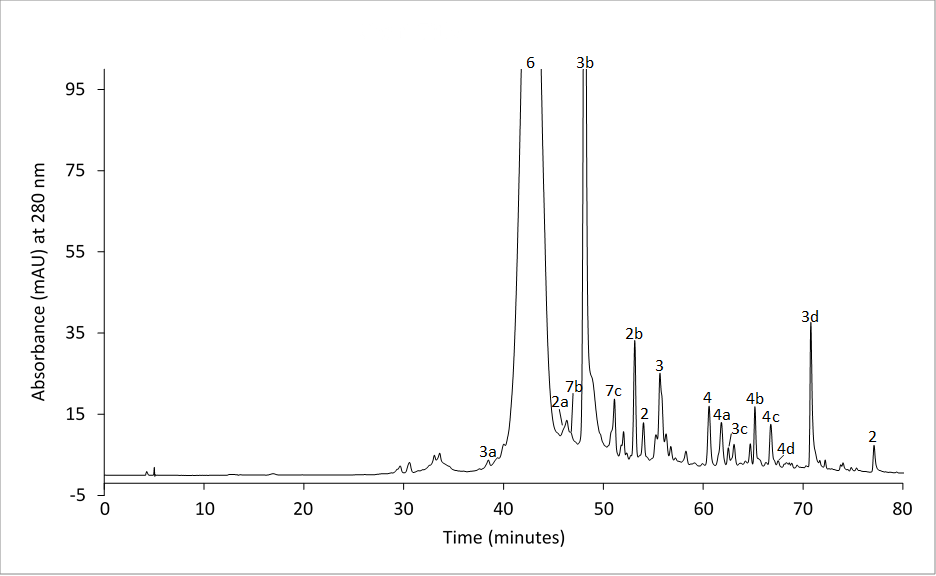

0

**B**

**A**

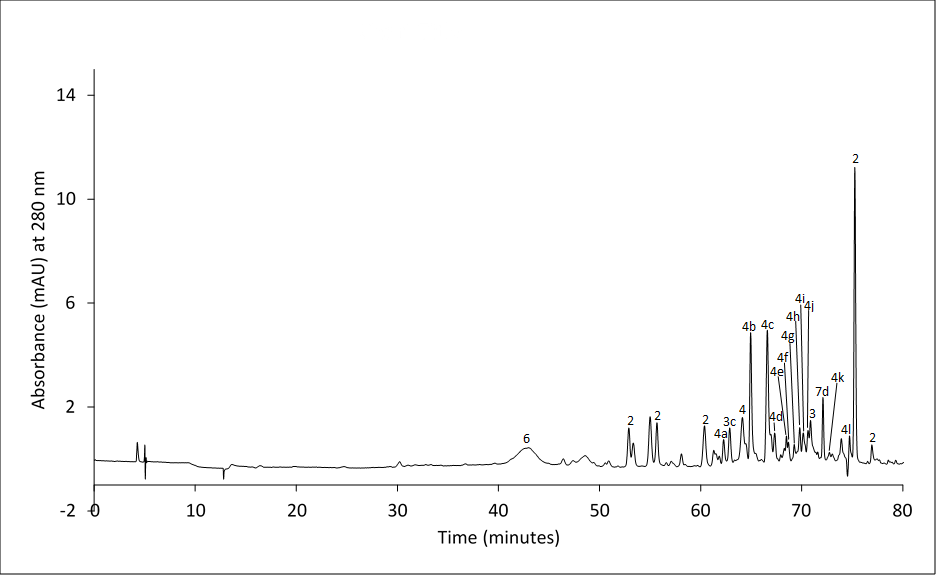

0

**C**

**Fig. S1B.** HPLC-PDA chromatograms of Tundra variety 40% ethanol (A), 70% ethanol (B), and 100% ethanol (C) fractions with peaks identified by HPLC-MS/MS and HPLC-PDA. Peak phenolic subclass assignments: 1. hydroxybenzoic acids; 2. flavanols; 3. hydroxycinnamic acids; 4. flavonols; 5. flavanones; 6. anthocyanins; and 7. other flavonoids. Peak identities: 2a. catechin^1^; 2b. epicatechin^1^; 3a. 3-*O*-caffeoylquinic acid^1^; 3b. 5-*O*-caffeoylquinic acid^1^; 3c. ferulic acid^1^; 3d. dicaffeoylquinic acid; 4a. quercetin-vicianoside; 4b. quercetin-3-*O-*rutinoside^1^; 4c. quercetin-3-*O*-glucoside^1^; 4d. quercetin-3-*O*-galactoside^1^; 4e. quercetin-pentoside; 4f. kaempferol-rutinoside; 4g. isorhamnetin-3-*O-*rutinoside^1^; 4h. kaempferol-hexoside; 4i. isorhamnetin-3-*O-*glucoside^1^; 4j. quercetin-hexoside-acetate; 4k. quercetin-hexoside; 4l. isorhamnetin-acetyl-hexoside; 7a. taxifolin-dihexoside; 7b. luteolin-hexoside; 7c. taxifolin-hexoside; 7d. phloridzin^1^. ^1^confirmed using standards

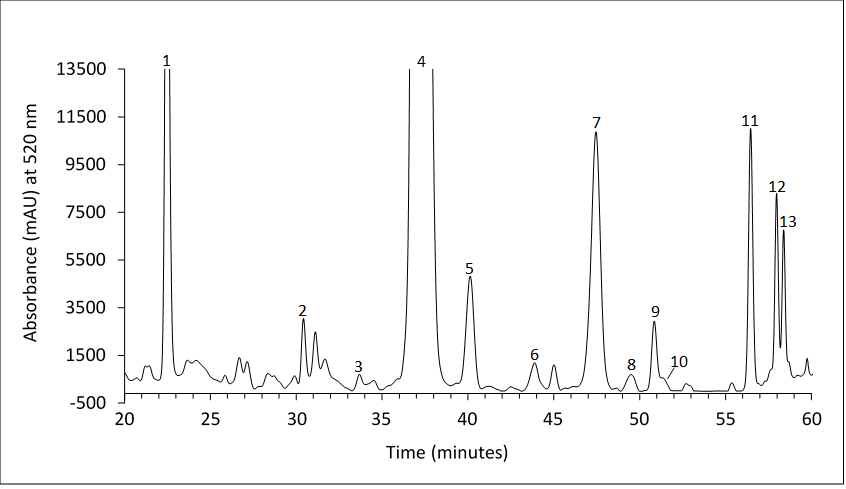

**Fig. S1C.** Representative HPLC-PDA chromatogram of the anthocyanins present in the Tundra 70% ethanol fraction. Peak assignments: 1. cyanidin-3,5-*O*-diglucoside^1^; 2. peonidin-dihexoside; 3. cyanidin-3-*O*-galactoside^1^; 4. cyanidin-3-*O*-glucoside^1^; 5. cyanidin-3-*O*-rutinoside^1^; 6. pelargonidin-3-*O*-glucoside^1^; 7. peonidin-3-*O*-glucoside^1^; 8. peonidin*-*rutinoside; 9. delphinidin-sambubioside; 10. cyanidin-3-*O*-xyloside^1^; 11. delphinidin-rutinoside; 12. delphinidin-hexoside; and 13. peonidin-pentoside. ^1^confirmed using standards.
